## Supplemental Materials for "XPRESSyourself: Enhancing, Standardizing, and Automating Ribosome Profiling Computational Analyses Yields Improved Insight into Data"

**Berg, et. al.**

#### Benchmarking Against TCGA Data

To further validate the design, reliability, and versatility of the XPRESSpipe pipeline, we processed raw TCGA sequence data using XPRESSpipe and compared the output count values to those publicly available through TCGA [1]. Spearman $\rho$ values for the selected samples ranged from 0.979-0.980 when pseudogenes were excluded (Figure 1), indicating XPRESSpipe performs with similar accuracy to the TCGA RNA-Seq processing standards.

The differences in reported counts can be accounted for by a couple of key differences. For instance, the XPRESSpipe-processed files are aligned to the *Homo sapiens* GRChv98 reference transcriptome, while the original count data are aligned to the GRChv79 reference transcriptome. The use of a different transcriptome reference can result in variance in the final quantified data for several genes (Figure 2) as significant advances have been made in our understanding of transcribed regions of the human genome between versions.

Another source of dissimilarity in data processing appears to arise if an Ensembl canonical transcripts-only reference is used during quantification. TCGA-processed data used an unmodified transcriptome reference file (all transcripts); therefore, the use of this modified (Ensembl canonical transcripts only) GTF will produce varied quantification for some genes as quantifications are constrained to a single transcript version of a given gene and a read will not be quantified if mapping to an exon not used by the canonical transcript. Even using XPRESSpipe settings closest to the TCGA pipeline and using the same genome and transcriptome version resulted in some variation (Figure 2, plot enclosed in maroon). By performing a more detailed analysis of these differences, it is clear that virtually all genes exhibiting variance between the processing methods are pseudogenes, with the TCGA pipeline accepting and quantifying more pseudogenes at the time of initial analysis of this dataset. This can be indicative of the difficulty surrounding the recognition of these reads as multi-mapping to both the original gene and pseudogene (Figure 3, 4, 5; interactive plots accompanying Figure 5 can be accessed at [2].

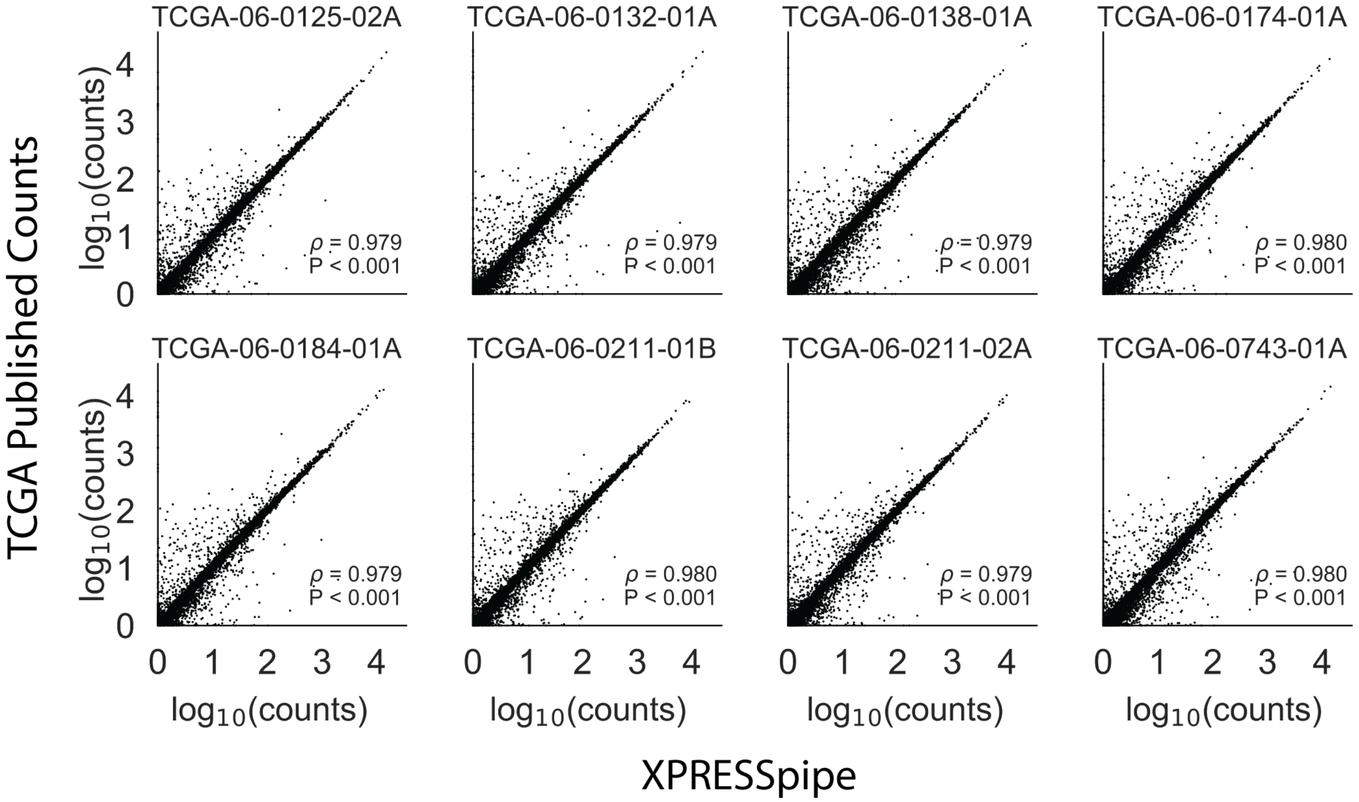


Figure 1: **Pipeline validation using publicly available TCGA count data.** Correlations were calculated between publicly available count data from TCGA samples and the count data processed by XPRESSpipe. Pseudogenes were excluded from the analysis. All reported $\rho$ values are Spearman correlation coefficients. XPRESSpipe-processed read alignments were quantified to *Homo sapiens* build CRCh38v98 using an unmodified GTF.

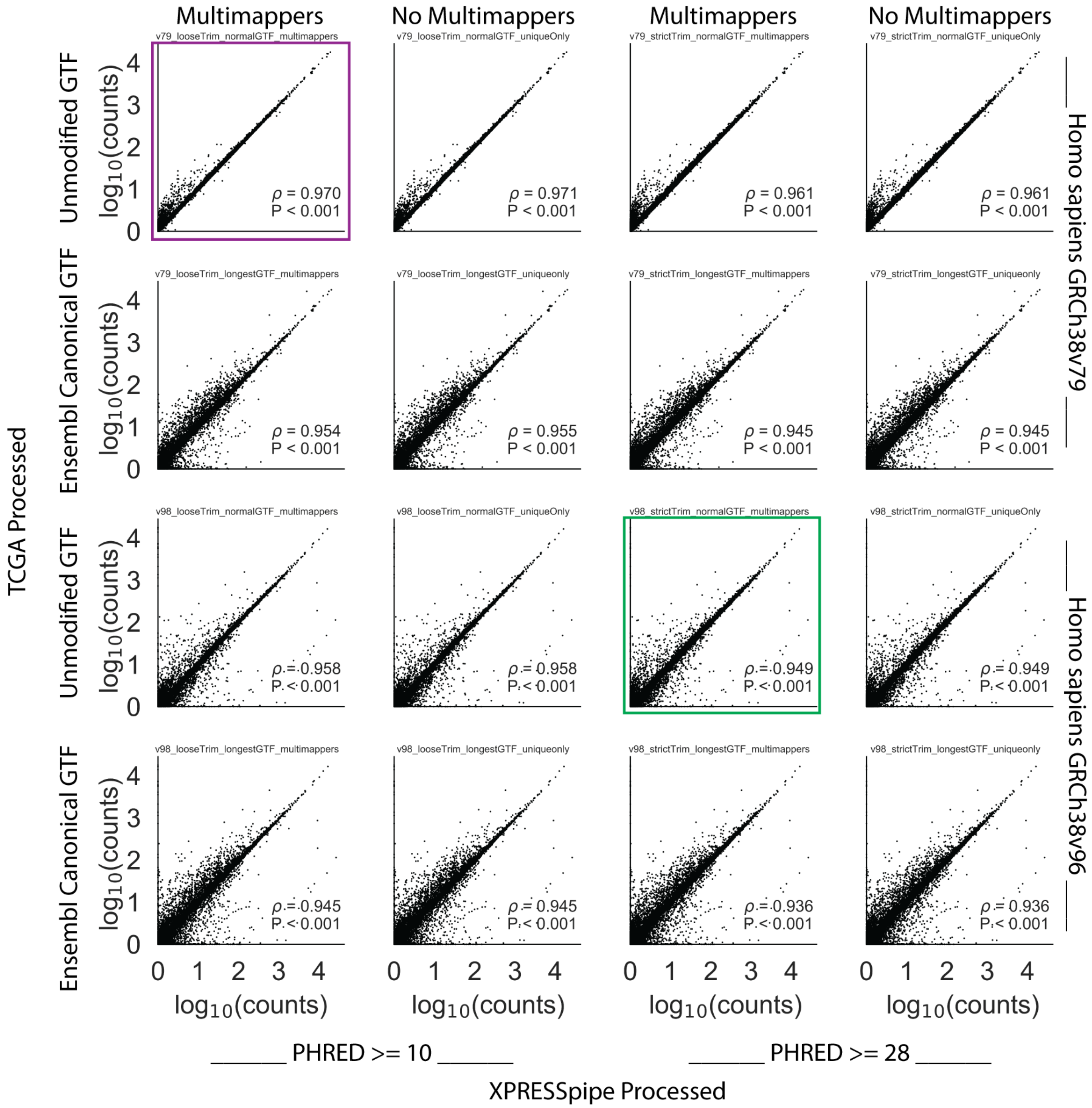


Figure 2: **Sample RNA-Seq count data compared between TCGA count data and various conformations of the XPRESSpipe pipeline.** An overview of how different conformations of the XPRESSpipe peRNAseq pipeline compared to the published TCGA sample TCGA-06-0132-01A count data. The x-axis data for the plot enclosed in maroon most closely mirrors the settings used in the published TCGA RNA-Seq pipeline. The x-axis data for the plot enclosed in green used XPRESSpipe default settings and the most current reference transcriptome at the time of writing. All $\rho$ values reported are Spearman correlation coefficients.

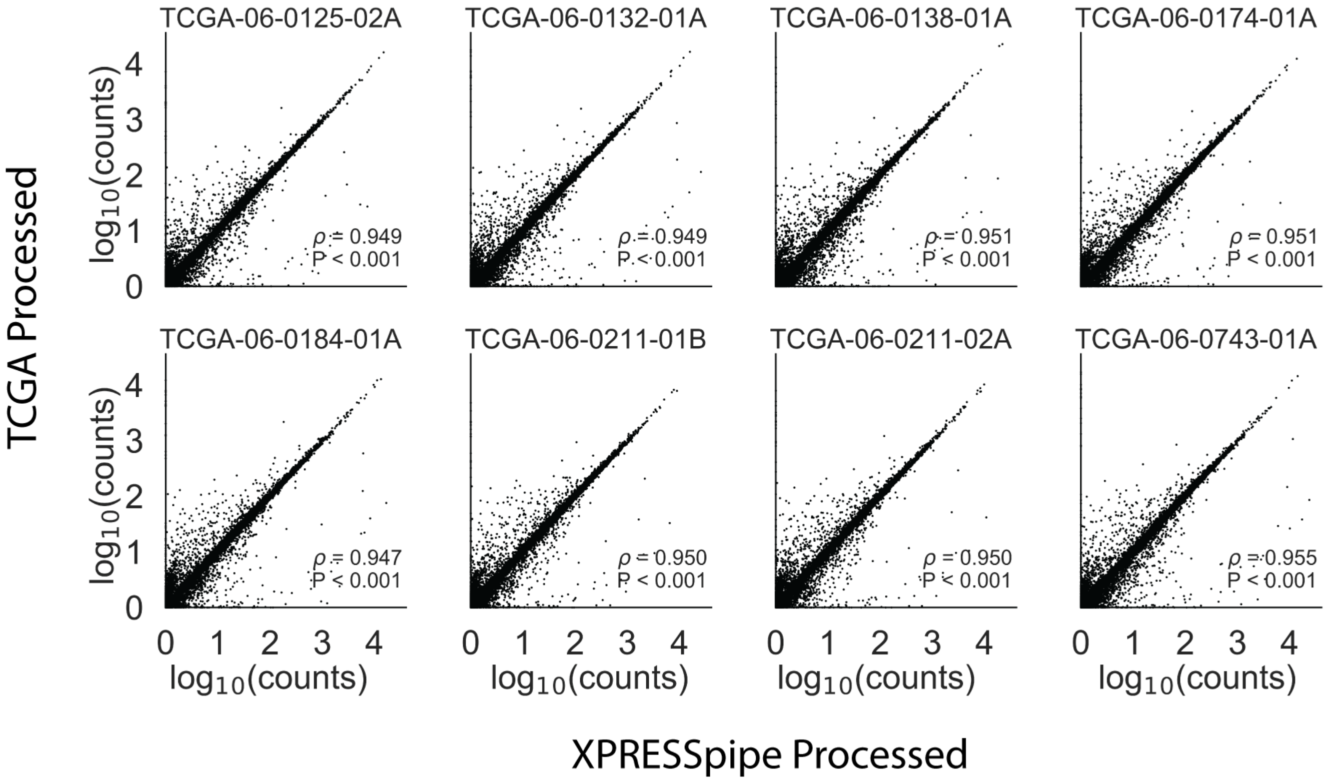


Figure 3: **Effect of pseudogene inclusion on comparability between processing regimes.** Spearman correlations between XPRESSpipe and TCGA-processed count data with pseudogene counts included. All $\rho$ values reported are Spearman correlation coefficients.

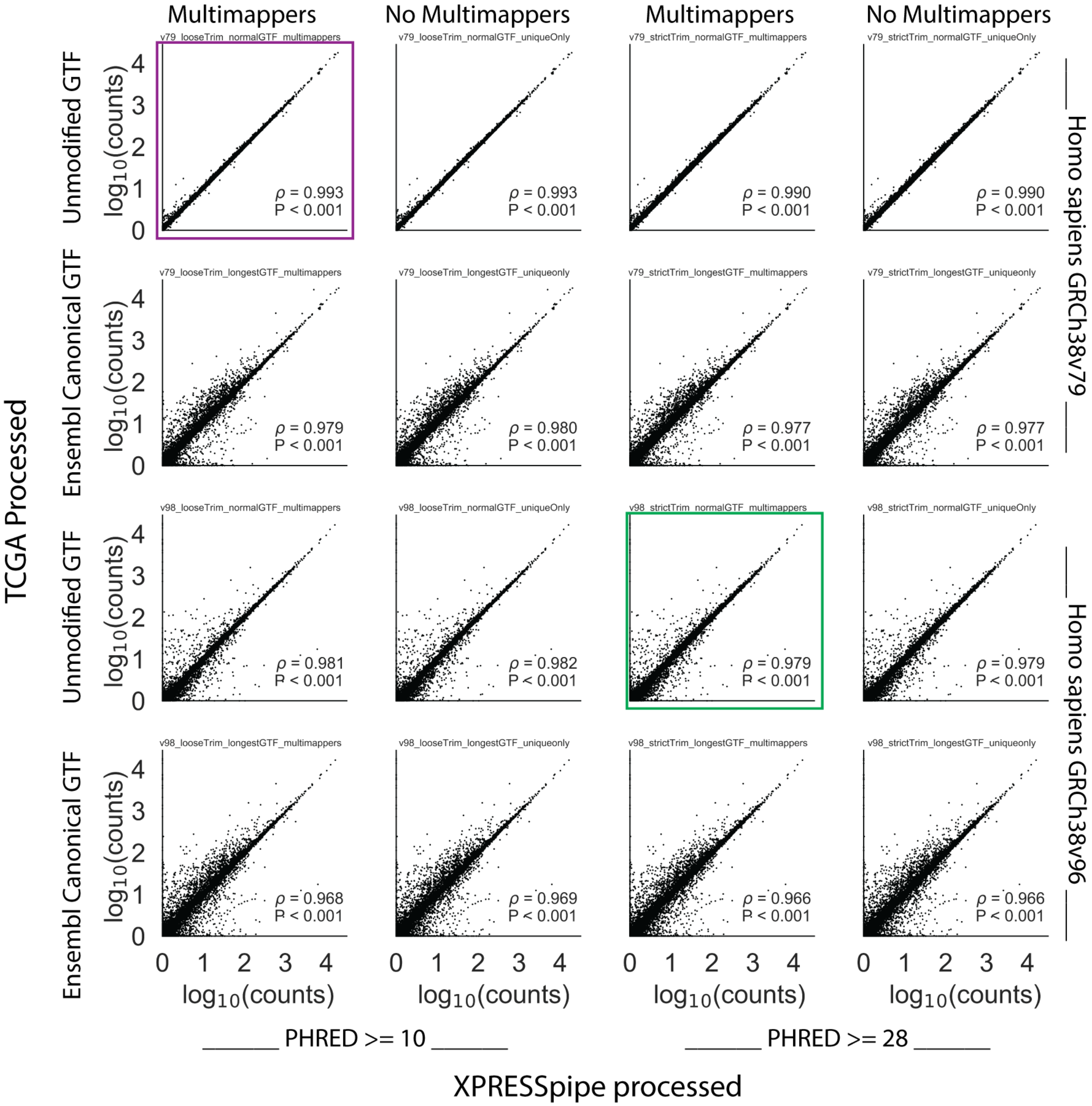


Figure 4: **Removal of pseudogenes counts improve comparability between analytical regimes.** An overview of how different conformations of the XPRESSpipe peRNAseq pipeline compared to the published TCGA sample TCGA-06-0132-01A count data with pseudogenes collapsed. The x-axis data for the plot enclosed in maroon most closely mirrors the settings used in the published TCGA RNA-Seq pipeline. The x-axis data for the plot enclosed in green used XPRESSpipe default settings and a current reference transcriptome. All $\rho$ values reported are Spearman correlation coefficients.

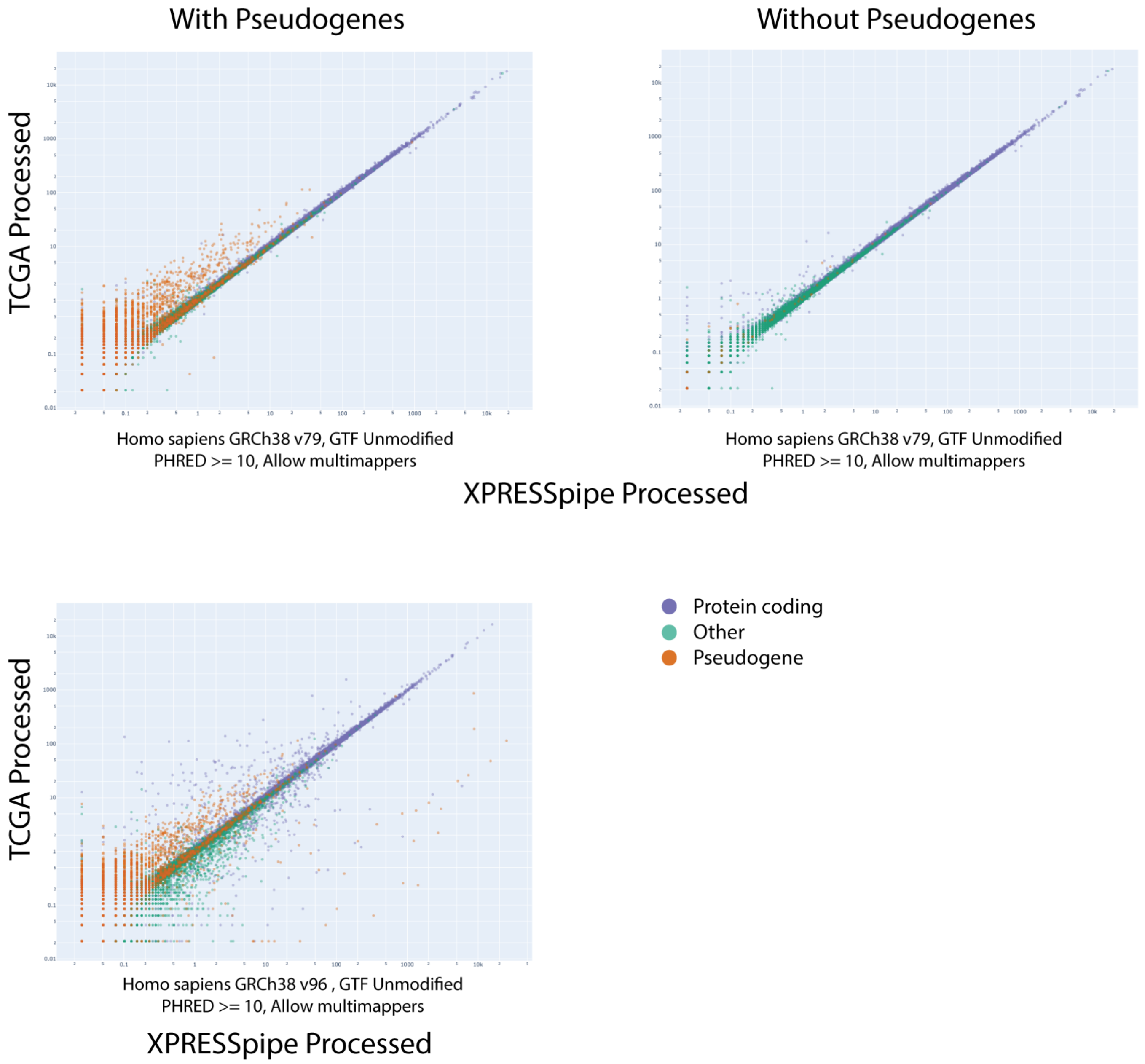


Figure 5: **Pseudogenes counts are over-represented in TCGA-processed data.** An overview of gene-type distributions between transcriptome reference versions. The plots above used GRCh38v79 and the bottom plot used GRCh38v98. Purple points, protein-coding genes. Orange points, pseudogenes. Green points, other gene records. All plots represent sample TCGA-06-0132-01A and were processed the same way except for transcriptome reference used during read quantification.

### Materials and Methods

Methods described in this manuscript apply to the software packages at the time of writing. To obtain the most current methods, please refer to the documentation or source code for a given module.

#### Software Dependencies

A list of dependencies required for XPRESSpipe at the time of writing is listed in Table 1. Dependencies for XPRESSplot at the time of writing are listed in Table 2. These are subject to change as these tools continue to evolve.

Table 1: **Summary of dependency software, accession location, and purpose in the XPRESSpipe package.**

| Package | Purpose | Reference |
|---|---|---|
| Python | Primary language | |
| R | Language used for some statistical modules | |
| Julia | Language used for file parsing features | |
| fastp | Read pre-processing | [3] |
| STAR | Reference curation and read alignment | [4] |
| samtools | Alignment file manipulation | [5] |
| bedtools | Alignment file manipulation | [6] |
| Cufflinks | Read quantification (primary) | [7] |
| HTSeq | Read quantification | [8] |
| FastQC | Quality Control | [9] |
| MultiQC | Quality Control | [10] |
| Pandas | Data manipulation | [11] |
| NumPy | Data manipulation | [12, 13] |
| SciPy | Data manipulation | [14] |
| scikit-learn | Data manipulation | [15] |
| Matplotlib | Plotting | [16] |
| XPRESSplot | Normalization and matrix manipulation | This paper |
| GenomicAlignments | BAM file processing | [17] |
| GenomicFeatures | GTF file processing | [17] |
| dupRadar | Perform library complexity calculations | [18] |
| riboWaltz | Perform p-site offset calculations | [19] |
| DESeq2 | Perform differential expression analysis | [20] |

#### Installation

XPRESSyourself suite packages can be easily installed following directions and/or walkthrough videos provided in the documentation and READMEs [24–26]. Once installed, `xpresspipe -h` can be run to explore sub-modules and their respective parameters.

XPRESSyourself software packages are hosted at https://github.com/XPRESSyourself. Current or past versions

Table 2: **Summary of dependency software, accession location, and purpose in the XPRESSplot package.**

| Package | Purpose | Reference |
|---|---|---|
| Python | Primary language | |
| R | Language used for some statistical modules | |
| Pandas | Data manipulation | [11] |
| NumPy | Data manipulation | [12, 13] |
| SciPy | Data manipulation | [14] |
| Matplotlib | Plotting | [16] |
| Seaborn | Plotting | [21] |
| Plotly | Interactive plotting | [22] |
| scikit-learn | Data manipulation | [15] |
| SVA | Perform batch correction for known effects | [23] |

of XPRESSyourself packages can be found at their respective repositories under the XPRESSyourself project. For simplest installation, we recommend using the Anaconda package manager [27]. Using the method below, you will need to activate the environment every time you wish to use XPRESSyourself software. Download XPRESSpipe's source code, set up the Anaconda environment, and install XPRESSpipe and XPRESSplot, as below (providing the version number you would like to download):

Listing 1: XPRESSyourself installation

```
$ curl -OL https://github.com/XPRESSyourself/XPRESSpipe/archive/XPRESSpipe-v0.3.0.tar.gz
$ tar xvzf XPRESSpipe-v0.3.0.tar.gz
$ cd XPRESSpipe-v0.3.0
$ conda env create --name xpresspipe -f requirements.yml
$ conda activate xpresspipe
$ python setup.py install
```

#### Running XPRESSpipe

Before processing data, the appropriate reference files must be curated. Download the FASTA and GTF files for the organism of interest (we recommend sourcing these from Ensembl [28]), change the GTF file name to `transcripts.gtf`, and run the code below. This will generate the appropriate STAR index files, as well as a GTF file with protein-coding transcripts only and the $5'$- and $3'$- end of each transcript truncated for ribosome profiling data. More examples of how to curate reference files can be found in the documentation [25], or one can use the `xpresspipe build` module.

Listing 2: curateReference example

```
$ xpresspipe curateReference \
           -o /path/to/reference/ \
           -f /path/to/reference/fasta_genome/ \
           -g /path/to/reference/transcripts.gtf \
           --protein_coding \
           --truncate \
           --truncate_5prime 45 \
```

```
        --truncate_3prime 15 \
        --sjdbOverhang 49 \
        --max_processors None
```

To process data, a simplistic example is provided below, but additional parameter changes may be necessary. Refer to the help menu (`xpresspipe -h`) or the documentation [25] for more information and examples of how to tune these parameters, or one can use the `xpresspipe build` module.

Listing 3: riboseq pipeline example

```
$ xpresspipe riboseq \
            -i /path/to/input_dir \
            -o /path/to/output_dir \
            -r /path/to/curated_reference \
            --gtf /path/to/transcripts_CT.gtf \
            -e isrib_dataset \
            -a CTGTAGGCACCATCAAT \
            --sjdbOverhang 49
```

Further analyses, such as additional gene coverage profiles, differential expression analysis, and batch normalization, can be performed. Details on how to perform these steps can be accessed from the help menu (`xpresspipe -h`) or the documentation [25].

For nearly all sub-modules, log files are written to the provided output directory to summarize provided user parameters, track performance, and report errors. An additional log file is written summarizing the versions of the different dependency software used during the execution of the pipeline or sub-module. Users are encouraged to provide these files as documentation when presenting XPRESSpipe-processed data.

#### Running XPRESSplot

Installation is handled during installation of XPRESSpipe, but can be installed independently:

Listing 4: XPRESSplot install

```
$ pip install xpressplot
<or>
$ conda install -c bioconda xpressplot
```

Generally, two inputs are required for all functions within XPRESSplot:

1. **Expression Matrix**: It is assumed that the input data matrix = *i* * *j* where *i* (rows) are genes or other analytes and *j* (columns) are samples.

2. **Metadata Table**: It is assumed that the metadata table is a two-column, header-less data matrix where column

0 is the sample ID (as specified in *j* column names of the expression matrix) and column 1 is the sample group (for example, genotype or treatment group).

More information on correct usage of all xpressplot functionalities can be found in the documentation [26].

**GTF Modification**

To parallelize GTF modification, a GTF file is split into approximately proportional chunks equal to the specified number of threads. To avoid an incomplete gene record being included in a chunk and being inappropriately processed, a given chunk endpoint is determined by calculating the size of the GTF, dividing by the number of available cores, and advancing to that endpoint, then advancing the endpoint line by line until the last line of the gene record encountered. This is performed for each subsequent chunk. If creating the last chunk, the end of the chunk is the last line of the GTF record. These steps are taken to ensure all transcripts are considered for a given transcript.

Ensembl canonical transcripts are determined according to the Ensembl glossary definition of a canonical transcript [29]. For cases where a tie exists between equal priority transcripts, the longest is chosen. When there are multiple transcripts that tie for equal priority and longest length, the first listed record is retained. Exon or CDS lengths are calculated by taking the sum of each exon or CDS, not including intron or other space in the calculation.

Protein-coding records are retained by performing a simple string search for the “protein_coding" annotation in the attribute column of a GTF file.

Truncation of records is performed by identifying the $5'$- and $3'$- end of each CDS and modifying the given coordinates to reflect the given truncation amounts. Suggested truncation amounts are 45 nt from the $5'$- end and 15 nt from the $3'$- end, both of which are set as the default truncation amount parameters for the function and do not need to be modified unless the user desires [30]. As a given CDS portion of a given exon may be less than a truncation amount, the function will perform a strand-aware recursive search CDS by CDS per transcript until the full truncation amount has been fully removed from each end. Any record smaller than the sum of the $5'$- and $3'$- truncation amounts is removed entirely from the output file. We include a table exploring the necessity of this truncation paradigm in Table 3.

Table 3: **Survey of GTF $5'$ CDS records requiring recursive truncation across selected model organisms.** Statistics were compiled using Ensembl v97 for most cases, with the exception of *Arabidopsis thaliana*, which used v44. # represents “number of". CDS represents coding sequence of gene.

| Organism | # CDSs requiring recursive truncation | # of transcripts total in organism | Percentage of recursive-requiring CDSs to total transcripts |
|---|---|---|---|
| *Arabidopsis thaliana* | 4,428 | 54,013 | 08.20% |
| *Caenorhabditis elegans* | 5,048 | 61,451 | 08.21% |
| *Danio rerio* | 8,987 | 59,876 | 15.01% |
| *Drosophila melanogaster* | 4,262 | 34,793 | 12.25% |
| *Homo sapiens* | 21,718 | 226,788 | 09.58% |
| *Mus musculus* | 12,779 | 142,333 | 08.98% |
| *Rattus norvegicus* | 5,002 | 41,078 | 12.18% |
| *Saccharomyces cerevisiae* | 160 | 7,127 | 02.24% |
| *Xenopus tropicalis* | 3,854 | 24,197 | 15.93% |

#### Flattened GTF Records

Flattened transcriptome references are created via a modified version of the annotation curation module available in riboWaltz [19]. Vectorized expressions in Pandas [11] are performed to quickly parse out pertinent meta-information for each transcript for the given analysis. Intermediate files are created for retrieval by each process when parallelizing analysis of each alignment file. This allows for fast processing of each BAM file, where the bottleneck in speed arises from the decompression and import of the binary alignment data (BAM file). Flat files are automatically destroyed after sub-module completion.

#### Normalization

Equations 1-4 reflect the design of the normalization functions within XPRESSplot, where *g* is gene *n*, *ge* is cumulative exon space for gene *n*, *r* is total reads, *f* is total fragments, and *l* is length.

$$RPM_{\text{g}} = \frac{1e6 \cdot r_{ge}}{\sum_{g=1}^{n} r_{ge}} \tag{1}$$

$$RPKM_{\text{g}} = \frac{1e9 \cdot r_{ge}}{(\sum_{g=1}^{n} r_{ge}) \cdot l_{ge}} \tag{2}$$

$$FPKM_{\text{g}} = \frac{1e9 \cdot f_{ge}}{(\sum_{g=1}^{n} f_{ge}) \cdot l_{ge}} \tag{3}$$

$$TPM_{\text{g}} = \frac{1e6 \cdot r_{ge}}{(\sum_{g=1}^{n} (\frac{1e3 \cdot r_{ge}}{l_{ge}})) \cdot l_{ge}} \tag{4}$$

#### Quality Control Summary Plotting

Summary plots are created using Pandas [11] and Matplotlib [16]. Kernel density plots for library complexity analyses are created using NumPy [12, 13] and SciPy's `gaussian_kde` function [14].

#### Metagene Estimation

Metagene calculations are performed by determining the meta-transcript coordinate *M* for each read alignment within a transcriptome-aligned BAM file (automatically output by STAR within XPRESSpipe). Let $L_e$ be the first mapped position of the read (strand agnostic and in reference to exon space to the $5'$-end) and *r* be the length of the mapped read. Let $\ell_e$ be the cumulative length of all exons for the given transcript. The subscripted *e* indicates the coordinate is relative to the exon space (intronic ranges within a transcript do not contribute to total space calculation). Extreme outliers (i.e., top and bottom 0.5% of transcripts ordered by their read abundances) are removed from analysis as they will inappropriately skew the meta-profile for the majority (99%) of transcripts. Similar methods are employed for metagene calculation across CDSs, where metagene coordinates are corrected for non-coding space of an exon. Required inputs are a transcriptome-aligned BAM file and a GTF reference file, which is flattened for downstream processing. For each mapped coordinate, the metagene position is calculated as:

$$M = \frac{(L_{\mathrm{e}} + \frac{1}{2}r) \cdot 100}{\ell_e} \tag{5}$$

#### Gene Coverage Plotting

Gene coverage calculations are performed by determining the exon space of the gene of interest and mapping any read for a given sample to this space. Each nucleotide of a read that maps to a nucleotide within these exon regions is counted. During plotting, a rolling window of 20 nucleotides is used to smoothen the plotted coordinates' read coverage. Required inputs are a transcriptome-aligned BAM file (as output by STAR within XPRESSpipe) and a GTF reference file, which is then curated into its longest-transcript, protein-coding-only flattened form, as discussed above. If a longest-transcript, protein-coding-only modified GTF has already been curated, this can alternatively be provided as input, with which the module will flatten (file suffix must be `LC.gtf`).

#### Periodicity

Ribosome p-site periodicity is calculated using riboWaltz [19]. Required inputs are the path to a directory containing transcriptome-aligned BAM files (as output by STAR within XPRESSpipe) and the path and file name of the appropriate unmodified GTF.

#### rRNA Probe

`rrnaProbe` works on a directory containing FastQC [9] zip compressed files to detect over-represented sequences for each sample. These sequences are then collated to create consensus fragments. One caveat of FastQC is that it collates on exact matching strings, but these strings, or sequences, can be 1 nt steps from each other and a single rRNA probe could be used to effectively pull out all these sequences. To handle this situation, XPRESSpipe will combine these near matches. A rank-ordered list of over-represented fragments within the appropriate length range to target for depletion is then output. A BLAST [31] search on a consensus sequence intended for probe usage can then be performed to verify the fragment maps to an rRNA sequence and is thus suitable for rRNA depletion.

#### Confidence Interval Plotting

Confidence intervals within PCA scatterplots generated by XRESSplot are calculated as follows:

1. Compute the covariance of the two principal component arrays, $x$ and $y$ using the `numpy.cov()` function.
2. Compute the eigenvalues and normalized eigenvectors of the covariance matrix using the `numpy.linalg.eig()` function.
3. Compute the $\theta$ of the normalized eigenvectors using the `numpy.arctan2()` function and converting the output from radians to degrees using `numpy.deg()`.
4. Compute the $\lambda$ of the eigenvalues by taking the square root of the eigenvalues.
5. Plot the confidence intervals over the scatter plot: The center point of the confidence interval is determined from the means of the $x$ and $y$ arrays. The angle is set equal to $\theta$. The width of the confidence interval is calculated by

$$w = \lambda_x \cdot ci \cdot 2$$

where *ci* is equal to the corresponding confidence level (i.e., 68% = 1, 95% = 2, 99% = 3). The height is similarly computed by

$$h = \lambda_y \cdot ci \cdot 2$$

**Ribosome Profiling Data Analysis**

Raw data were obtained from GEO (GSE65778). Reference files were taken from Ensembl Human build GRCh38 version 98. Read alignments were quantified using an XPRESSpipe-modified GTF file that contained only protein-coding records and the $5'$- ends of each CDS truncated by 45 nucleotides and the $3'$- ends truncated by 15 nucleotides. All associated figures and analyses can be reproduced using the associated scripts found at [2].

Only gene names in common between the original data file and XPRESSpipe output were used for the method comparisons. Correlation between methods or replicates were calculated using a Spearman rank correlation coefficient, performed using the `scipy.stats.spearmanr()` function [32]. Pearson correlation coefficients were calculated using $\log_{10}$(rpm(counts) + 1) transformed data and the `scipy.stats.pearsonr()` function. Violin plots were generated using Seaborn [21]. RNA-Seq samples were thresholded at 10 counts.

Differential expression analyses were performed using all genes, but with a minimum count of 25 or greater per gene across samples, as recommended by the DESeq2 documentation [20]. Differential expression for ribo-seq and RNA-Seq was performed as detailed in the associated scripts [2]. For these analyses, the design formula was such that comparisons were designed as “treated" factor level over “untreated" factor level. Differential expression of translation efficiencies between conditions used the additional incorporation of the “ribosome footprint" factor level over “RNA-Seq" factor level in the design formula [20,30,33]. Adjusted p-values (FDRs) in the associated figures were calculated from the differential expression of the translation efficiencies of each gene for a given condition. Those passing an adjusted p-value (FDR) threshold of less than or equal to 0.1 are highlighted in black. Venn diagrams were produced using matplotlib-venn [34]. Differentially expressed genes were for these diagrams were obtained by using the same thresholds as in the original study (RPF fold change $\geq$ 1 or $\leq$ -1, FDR $<$ 0.1). FDR values used here were obtained from the two-factor design formula used in DESeq2 [20]. Differential expression lists from the original study were obtained from the manuscript’s supplemental files [33].

Intron-agnostic gene coverage profiles were generated using XPRESSpipe’s `geneCoverage` module. Comparison plots were generated using IGV [35]. Interactive scatter plots were generated using Plotly Express [22].

#### TCGA Data Analysis

Raw data and processed TCGA count data was obtained from the TCGA Portal [1] via dbGap controlled access [36]. Raw data were processed on a protected high-performance computing environment. Correlations between methods or replicates were calculated using a Spearman rank correlation coefficient, performed using the `scipy.stats.spearman()` function [32]. Interactive scatter plots were generated using Plotly Express [22]. The associated scripts can be accessed at [2].

#### Cost Analysis

Cost analysis was performed by assessing run logs from the high-performance computing cluster and using published AWS prices [37,38] (accessed 29 October 2019) to calculate the relative cost for a comparable run. Pricing assumed the user was in Salt Lake City, Utah, USA, using the US East (Ohio) AWS server. Sub-module statistics were compiled by running each step of the pipeline with the command `/usr/bin/time -v`.

### References


[1] The Cancer Genome Atlas. https://portal.gdc.cancer.gov.

[2] Manuscript code. https://github.com/XPRESSyourself/xpressyourself_manuscript/tree/master/supplemental_files. Available from: https://doi.org/DOI: 10.5281/zenodo.3337599.

[12] T. Oliphant. *A guide to NumPy* (Trelgol Publishing, USA, 2006).

[22] P. T. Inc.. Collaborative data science (2015).

[23] J. Leek, W. Johnson, H. Parker, A. Jaffe, J. Storey. The sva package for removing batch effects and other unwanted variation in high-throughput experiments. *Bioinformatics* **28** (2012). Available from: https://doi.org/10.1093/bioinformatics/bts034.

[24] XPRESSyourself. `https://github.com/XPRESSyourself/`.

[25] XPRESSpipe documentation. `https://xpresspipe.readthedocs.io/en/latest/`.

[26] XPRESSplot documentation. `https://xpressplot.readthedocs.io/en/latest/`.

[27] Anaconda. Anaconda Software Distribution. `https://anaconda.com`.

[28] Ensembl. `https://uswest.ensembl.org`.

[29] Ensembl. `https://uswest.ensembl.org/Help/Glossary`.

[30] N. McGlincy, N. Ingolia. Transcriptome-wide measurement of translation by ribosome profiling. *Methods* **126**, 112 (2017). Available from: https://doi.org/10.1016/j.ymeth.2017.05.028.

[31] S. Altschul, W. Gish, W. Miller, E. Myers, D. Lipman. Basic local alignment search tool. *J Mol Biol.* **215**, 403 (1990).

[32] F. Liesecke, *et al.*. Ranking genome-wide correlation measurements improves microarray and RNA-seq based global and targeted co-expression networks. *Sci Rep* **8** (2018). Available from: https://doi.org/10.1038/s41598-018-29077-3.

[33] C. Sidrauski1, A. McGeachy, N. Ingolia, P. Walter. The small molecule ISRIB reverses the effects of eIF2ã phosphorylation on translation and stress granule assembly. *eLife* (2015). Available from: https://doi.org/10.7554/eLife.05033.

[34] K. Tretyakov. matplotlib-venn. `https://github.com/konstantint/matplotlib-venn`.

[35] J. Robinson, *et al.*. Integrative Genomics Viewer. *Nat Biotechnol* **29**, 24 (2011). Available from: https://doi.org/10.1038/nbt.1754.

[36] Database of Genotypes and Phenotypes. `https://www.ncbi.nlm.nih.gov/gap/`.

[37] Amazon Web Services EC2 Computing. `https://aws.amazon.com/ec2/pricing/on-demand/`. Accessed 29 October 2019.

[38] Amazon Web Services S3 Storage. `https://aws.amazon.com/s3/pricing/`. Accessed 29 October 2019.
